# Probing cytoskeletal modulation of passive and active intracellular dynamics using nanobody-functionalized quantum dots

**DOI:** 10.1101/089284

**Authors:** Eugene A. Katrukha, Marina Mikhaylova, Hugo X. van Brakel, Paul M. van Bergen en Henegouwen, Anna Akhmanova, Casper C. Hoogenraad, Lukas C. Kapitein

## Abstract

The cytoplasm is a highly complex and heterogeneous medium that is structured by the cytoskeleton. Cytoskeletal organization and dynamics are known to modulate cytoplasmic transport processes, but how local transport dynamics depends on the highly heterogeneous intracellular organization of F-actin and microtubules is poorly understood. Here we use a novel delivery and functionalization strategy to utilize quantum dots (QDs) as probes for transport dynamics in different sub-cellular environments. Rapid imaging of non-functionalized QDs revealed two populations with a 100-fold difference in diffusion constant. Depolymerization of actin increased the fast diffusing fraction, suggesting that slow QDs are trapped inside the actin network. When nanobody-functionalized QDs were targeted to different kinesin motor proteins and moved over microtubules, they did not experience strong actin-induced transverse displacements, as suggested previously. Only kinesin-1 bound QDs displayed subtle directional fluctuations, because the specific subset of stable microtubules used by this motor underwent prominent undulations. Using actin-targeting agents we found that the actin network suppresses most microtubule shape remodeling, rather than promoting them. These results demonstrate how the spatial and structural heterogeneity of the cytoskeleton imposes large variations in non-equilibrium intracellular dynamics.

## Introduction

Cells have a highly structured internal organization that depends on the cytoskeleton, a network of biopolymers, including F-actin and microtubules, that shapes the cell and enables active intracellular transport driven by a variety of motor proteins that can walk along F-actin (i.e. myosins) or microtubules (i.e. kinesins and dyneins). Intracellular transport processes can be categorized as either active or passive, depending on whether they are driven by chemical energy or by thermal excitation, respectively. While linear motion driven by molecular motors is clearly active, recent studies have argued that the apparent passive dynamics of intracellular particles is largely driven by the active contractile dynamics of the actomyosin network, rather than by thermal excitation^9-11^. Although one study suggested that even the apparent diffusion of individual proteins depends on this non-thermal mixing^10^, it has remained unclear for which particle sizes fluctuations of the viscoelastic actomyosin network would dominate over thermal diffusion. In addition, given that actin is enriched near the cortex, non-thermal contributions are likely to depend on the position within the cell, but how the highly heterogeneous intracellular organization of F-actin affects passive particle dynamics has remained largely unexplored.

The heterogeneous composition of the cytoplasm should also affect active point-to-point transport inside cells^12-14^. Recently, the intracellular behavior of custom-synthesized single-walled carbon nanotubes (SWNT) was probed over several orders of magnitude in time and space^11^. It was reported that their dynamics was strongly affected by the presence of an active viscoelastic cytoskeleton and that an active stirring mechanism induced large sideways fluctuations of SWNTs that were transported over microtubules by kinesin-1. SWNTs are non-isotropic rods and have a disperse length distribution that spans at least one order of magnitude (100-1000 nm), which will result in disperse hydrodynamic properties that could hamper proper analysis and interpretations. Although they are only 1 nm in diameter, their stiffness is comparable to that of actin filaments^11^. Given these lengths and stiffness, their dynamics is expected to be governed by the actin network, which has a characteristic mesh size of ~100 nm^15^. Importantly, earlier work reported that kinesin-1 moves on a specific subset of microtubules and, therefore, the generality of these observations has remained unclear^16^.

Here we use quantum dots (QDs) to examine how the actin cytoskeleton modulates active and passive transport in different subcellular zones. QDs are an attractive alternative for SWNTs and plastic beads, because they are widely available, can be tuned to emit in the visible range and have a monodisperse size distribution. Although QDs have been used previously to study intracellular dynamics, their widespread use in intracellular applications has so far been hampered by challenges in intracellular delivery and functionalization^17^. We combined adherent cell electroporation and nanobody-based functionalization to deliver QDs to the cytoplasm and analyzed the dynamics of both non-functionalized QDs and QDs that are targeted to different subsets of microtubules by binding to different kinesin family members.

We report that the diffusion of non-functionalized QDs was highly heterogeneous and dependent on the presence of F-actin. Most microtubule-associated QDs propelled by different types of kinesin did not experience strong (actin-induced) transverse fluctuations. Kinesin-1 bound QDs, however, displayed directional fluctuations, because the subset of modified microtubules used by this motor underwent more prominent undulations. However, this shape remodeling was not caused by active contractility of actomyosin network, but instead suppressed by it. These results demonstrate how the heterogeneity of the mammalian cytoskeleton imposes a large spatial variation in non-equilibrium cellular dynamics, which precludes straightforward application of physical approaches that model the cytoplasm as a viscoelastic homogeneous and isotropic medium.

## Results

### Rapid and slow diffusion of cytoplasmic quantum dots

As small size intracellular probes we used commercially available QD625-streptavidin conjugates with a measured Stokes radius of 14.4 nm (Supplementary Fig.1a). To explore the passive dynamics of QDs inside the cytoplasm, we used adherent cell electroporation for intracellular delivery of particles. After optimization of electroporation parameters (Supplementary Table 1), we observed that most cells routinely enclosed 10-100 QDs (Fig. 1a,b). Upon internalization, we observed a strong suppression of QDs blinking, manifested in the increase of the average duration of the on-state (Supplementary Fig.1c). This is likely the result of the mildly reducing environment of the cytoplasm, which is known to reduce QD blinking^18^.

**Figure 1:**
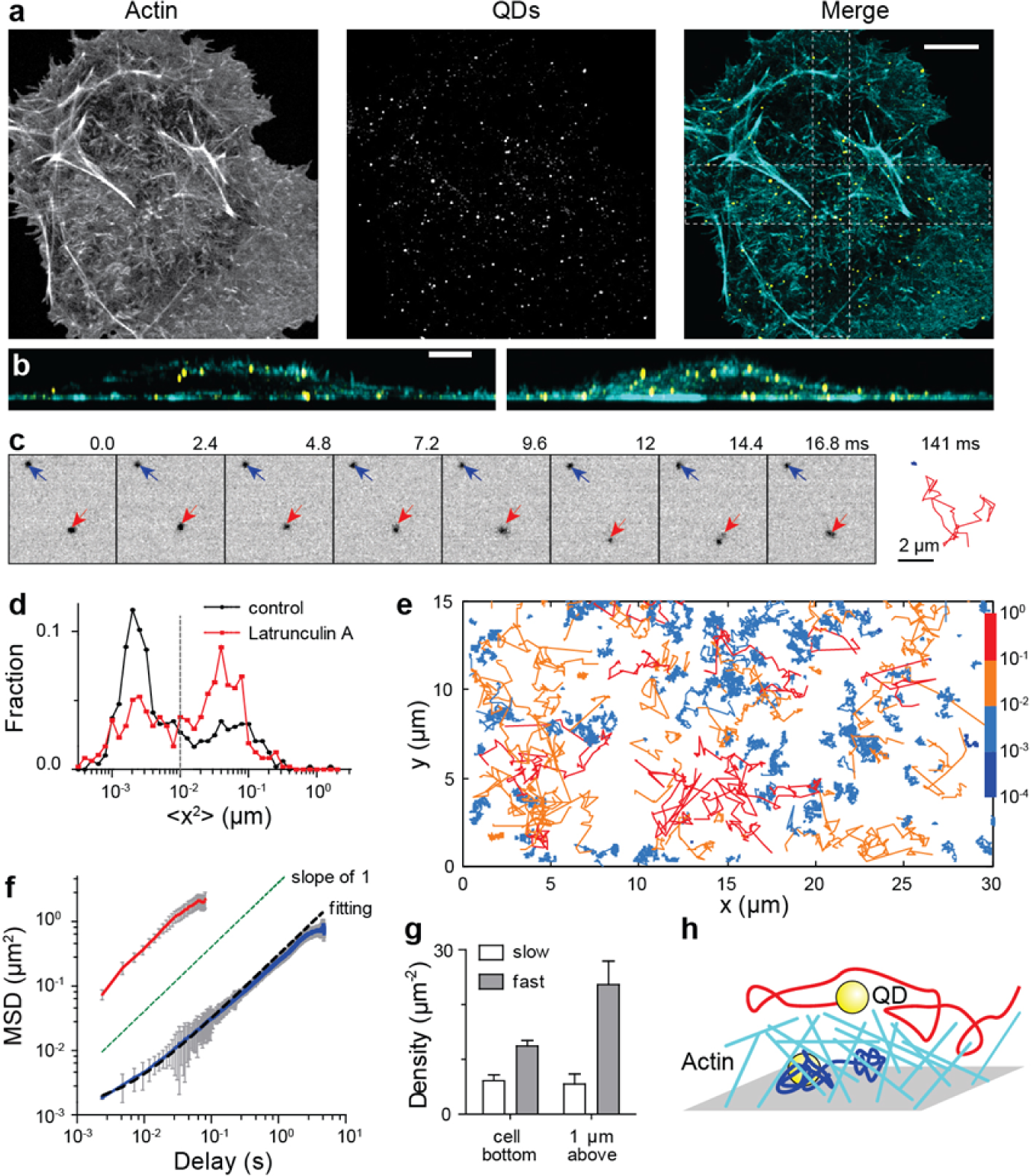
Non-isotropic diffusion of intracellular QDs. (a) COS-7 cell fixed 30 minutes after electroporation with QDs and stained with phalloidin. Maximum projection of a z-stack acquired with the spinning disk microscope. Scale bar 10 μm. (b) Lateral Y-Z (left) and X-Z maximum projection views of cross sections along the boxes depicted in **a**. Scale bar 5 μm. (c) Stills from a stream recording of GFP-actin expressing COS-7 cell electroporated with QDs. The interval between frames is 2.4 ms. Blue and red arrows indicate slow and fast cytosolic diffusion of QDs, respectively. The complete trajectories (66 frames, 156 ms) are depicted on the right panel. (d) Distribution of the mean square displacement at one frame (2.4 ms) delay for the trajectories of QDs in control (black, N=484, 10 cells) and latrunculin A treated cells (red, N=474, 13 cells). Dashed line marks the threshold used to separate trajectories of fast and slow diffusing particles in F. (e) Example trajectories of intracellular QDs, color-coded according to the corresponding value of mean square displacement at one frame (2.4 ms) delay. (f) Average MSD of slow (blue solid line, N=317) and fast (red solid line, N=167) fractions of QDs trajectories. Error bars represent SEM. Dashed line shows fit MSD(*τ*) = 4*Dτ* + *dx*^2^ where offset *dx*^2^=35 nm^2^ reflects squared average localization precision. (g) Density of fast and slow diffusing subpopulation of QDs in cells, imaged either near the coverslip or 1 μm above (N=6 cells per each condition). (h) Proposed origin of slow and fast diffusing subpopulations of QDs.

Rapid live-imaging with 2.4 ms intervals of QDs located within the cell’s lamella, followed by single-particle tracking and MSD analysis, revealed fast and slow diffusing subpopulations of QDs (Fig. 1c-e and Supplementary Movie 1), which appeared as two separate peaks in the distribution of the mean squared frame-by-frame displacement (Fig. 1d). For some QD we also observed transitions from slow to fast mobility regimes and back (Supplementary Fig.1b). The fraction of slowly diffusing QDs was strongly reduced after treatment with latrunculin A, an inhibitor of actin polymerization (Fig. 1d). The average MSD curves for the fast and slow fractions yielded diffusion coefficients of *D*_fast_ = 10.1 μm^2^ s^−1^ and *D*_slow_ = 0.06 μm^2^ s^−1^, respectively (Fig. 1f). Thus, rapid intracellular tracking of QDs revealed two classes of QD diffusion, the faster of which could be promoted by actin destabilization.

The increased diffusion constant upon actin depolymerization suggests that slowly diffusing QDs are trapped within the F-actin-rich cellular cortex at the inner face of the plasma membrane. Optical cross sections of cells labelled for F-actin show that F-actin density is high close to the cell membrane, bus strongly decreases at 1 μm depth into the cell (Fig.1b). We therefore also measured the density of fast and slow moving particles at 1 μm above the cell’s bottom cortex and compared it with the area close to a coverslip (Fig.1g). Indeed, the density of rapidly diffusing QD increased from 12.4 to 23.7 μm^−2^ when moving 1 μm deeper into the cell and the density of slowly diffusing QDs remained almost unchanged (6.1 to 5.5 μm^−2^). These results indicate that upon electroporation of adherent cells, most QDs are freely moving in the internal cytoplasm and suggest that slow diffusive QDs are trapped within the F-actin-rich cellular cortex at the inner face of the plasma membrane (Fig. 1h). In standard recordings with 100 ms intervals, rapid QD diffusion will not be detectable and observations will be biased towards slower diffusing QDs embedded in actin meshwork (Supplementary Movie 2).

### Probing specific motor proteins using nanobody-conjugated QDs reveals limited transverse fluctuations during microtubule-based runs

Previous work using single-walled carbon nanotubes reported that cargoes propelled by kinesin-1 experience significant sideways fluctuations (~0.5-1μm), which were ascribed to myosin-driven fluctuations of the actin cytoskeleton. The generality of these observations has remained unclear, because not all microtubules are embedded within F-actin and because kinesin-1 has been reported to move over a specific subset of stabilized microtubules^16^. To perform the same assay using probes of smaller size, we labeled QDs with biotinylated nanobodies (VHH_GFP_) against GFP. Nanobodies are promising candidates for QD functionalization because they are small, very stable, and easily produced ^19^ (Supplementary Fig. 1d-h). We delivered these nanobody-functionalized QDs into cells transfected with different constitutively active GFP-tagged motor proteins. COS-7 cells expressing kinesin-1 showed a peripheral accumulation of QDs-VHH_GFP_ (Fig. 2a-c), and live-imaging revealed numerous QDs running processively over the kinesin-1 decorated microtubules (Fig. 2c-d; Supplementary Movie 3). Similar results were obtained for kinesin-2 (KIF17-GFP-FRB), kinesin-3 (KIF1A-GFP-FRB) and kinesin-4 (KIF21B-GFP-FRB). The bright fluorescence of QDs allowed us to reach a localization precision of 3-5 nm (Supplementary Table 3), while recording the fast motion using continuous stream acquisition with 50 ms exposure without any noticeable photobleaching. All motor proteins tested in our QDs transport assay showed dynamic binding to the microtubules, i.e. processive movement interspersed with dissociation and cytosolic diffusion (Fig. 2d-h), and moved unidirectionally, suggesting that QDs are bound exclusively to the respective GFP-labeled kinesins^20^.

**Figure 2:**
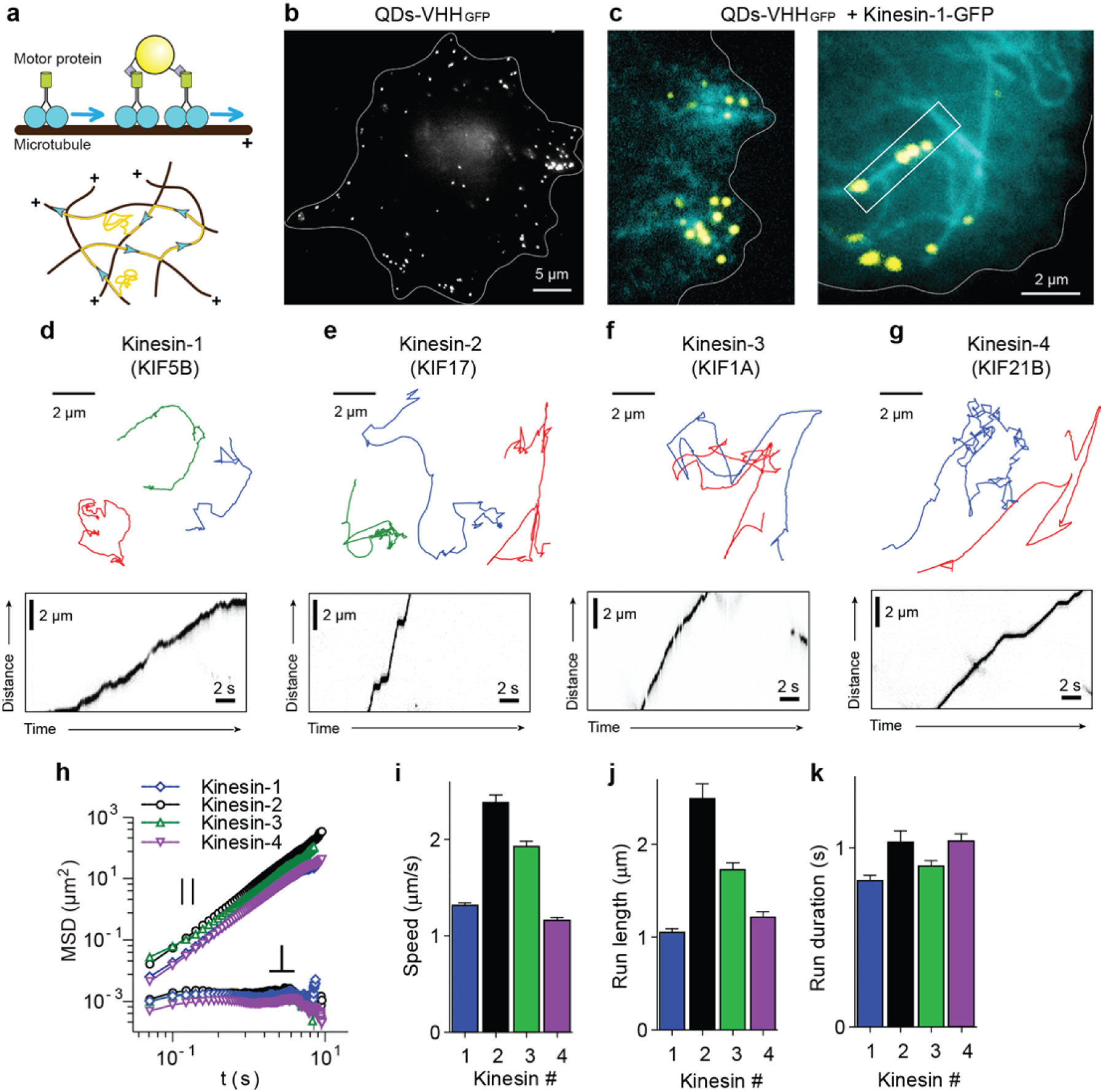
Probing specific motor proteins using nanobody-conjugated QDs reveals limited transverse fluctuations during microtubule-based runs. (a) Linkage of QDs to GFP-fused motor proteins through GFP nanobody (top) and expected movement of individual QDs-kinesin complexes along microtubules (bottom). (b) Distribution of electroporated QDs-VHH_GFP_ inside COS-7 cell expressing kinesin-1-GFP (cyan) and electroporated with QDs-VHH_GFP_ (yellow). White curve indicates cell outline. (c) QDs-VHH_GFP_ colocalize with microtubules decorated by kinesin-1 (KIF5B-GFP-FRB). Single frames from TIRFM stream recordings of COS-7 cell expressing kinesin-1-GFP (cyan / right) and electroporated with QDs-VHH_GFP_ (yellow). (**d-g**) Example trajectories (top row) and kymographs (bottom row) for QDs coupled to kinesins from different families. (**h**) Average MSD of longitudinal (top) and transverse (bottom) components of directed motion segments of different kinesins (n=104, 146, 151, 144 for Kinesin-1,2,3,4) decomposed using B-spline trajectory fitting with 1 μm control points distance. (**i-k**) Average speed **i**, run length **j** and run duration **k** of the individual motor runs for the different kinesins. Error bars represent S.E.M. See Supplementary Table 3 for numeric values.

To robustly extract the transient periods of directional movement from the trajectories, we developed a set of directional filtering techniques that was validated on artificial data (Supplementary Fig. 2a-f, and Supplementary Methods). The filtering was able to detect both strictly directional movements of motors with a smooth change of velocity’s angle and those containing abrupt stochastic displacements of underlying microtubules (Supplementary Fig. 2e,f). To characterize the transverse and longitudinal dynamics of directionally moving QDs relative to the microtubule axis, we estimated the position of the microtubule by fitting continuous non-periodic cubic B-splines to the QD coordinates (Supplementary Fig. 2d) and projecting the coordinates onto these curves. The flexibility of the spline curve is defined by the number of internal knots and control points placed along its arc length and the distance between such control points sets the length scale at which microtubules are considered straight (Supplementary Fig. 2g). We found that the behavior of the transverse component was heavily dependent on the degree of spline approximation (Supplementary Fig. 2g-j). When B-spline control points were positioned with 1 μm interval, the amplitude of transversal displacement diminished in comparison to 6 μm and the transverse MSD curve changed its behavior from diffusional to highly confined (Supplementary Fig. 2h-j).

For all further analyses, we chose the distance between control points equal to 1 μm to account for the highly bent microtubule shapes observed in live cells, which are relatively stable on the time scale of kinesin movement (~ 1 s, Fig. 2k). MSD analysis of trajectories of different kinesin family members revealed the same highly confined transverse behavior (Fig. 2h), whereas the longitudinal MSD increased quadratically with time delay *τ*, as expected for directional movement. For all directional movement episodes, we determined the average speed run length and run duration for the different kinesin family members (Fig. 2i-k). The speeds ranged from 1 μm s^−1^ for kinesin-1 to 2.5 μm s^−1^ for kinesin-2, similar to results obtained for non-cargo bound motors^16^. Interestingly, reported kinesin-1 speeds measured using SWNT were 3-fold lower, presumably due to the non-selective interactions with other cellular components^11^. These results demonstrate that, unlike reported previously^11^, most motor-driven particles do not undergo large sideways fluctuations.

### Kinesin-1 targeted microtubules are actively bent by longitudinal forces

As shown above, the underfitting of trajectories will affect the MSD analysis of the transverse dynamics. Conversely, overfitting could result in trajectories segments with unrealistically high curvatures that no longer reflect the underlying microtubule shape. To verify the consistency of our analysis, we built the distribution of trajectory curvatures for all four motors tested and found that the curvature of about 90% of arc length of runs is below 2 μm^−1^ (i.e. radius of >0.5 μm, Fig. 3a,b), meaning that the fraction of highly curved segments is low. Surprisingly, there was a clear difference between the curvature distributions of kinesin-1 and other kinesins (Fig. 3b,c). The processive runs of kinesin-1 were more curved, suggesting that this motor runs on a subset of microtubules that are more bent.

**Figure 3:**
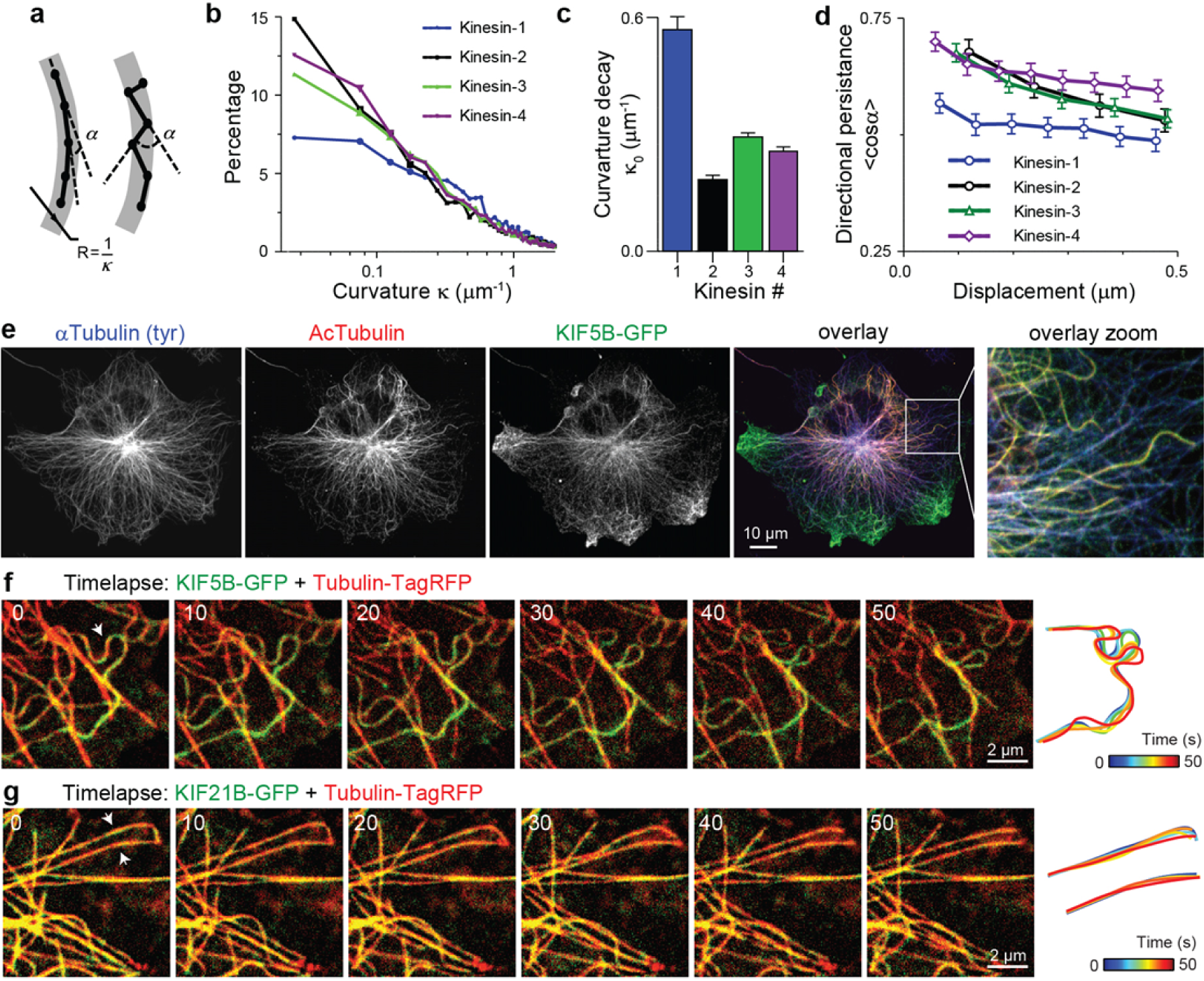
Kinesin-1 preferably walks on highly curved, acetylated microtubules that undergo bending deformations. (a) Example of two trajectories with the same radius of curvature R, but different average angle between the directions of consecutive displacements. (b) Distribution of local curvatures of directed motion segments in trajectories of QDs coupled to different kinesins. (c) Characteristic decay values from exponential fits to distributions in **b**. Error bars represent standard error of fitting. (d) Average directional persistence (cosine between consecutive displacements) as a function of displacement, assuming constant average speed of kinesins (n is the same as in Fig. 3H, see also Supplementary Fig. 4a). (e)COS-7 cell expressing KIF5B-GFP (green) stained for tyrosinated (blue) and acetylated (red) tubulin. (**f**),(**g**) Stills from a time lapse recording of a COS-7 cell expressing KIF5B-GFP (F) or KIF21B-GFP (G) (green) and Tubulin-TagRFP (red). Shapes of microtubules highlighted by white arrows are traced over time (sec) and color-coded as indicated.

The higher curvature of kinesin-1 targeted microtubules could be caused by active remodeling forces that are selective for these microtubules. Such shape remodeling would affect the smoothness of QD trajectories and we therefore examined the directional persistence of trajectories by calculating the correlation length of the angle between subsequent displacements. We found that the directional persistence was lower for kinesin-1 in comparison to other kinesins (Fig. 3d and Supplementary Fig. 3a), indicating that the amount of stochastic distortion during processive runs is higher. These results suggest that the microtubules targeted by kinesin-1 are fluctuating more than microtubules targeted by the other kinesins.

It has been reported that kinesin-1 moves preferentially on a subset of stable microtubules marked by certain post-translational modifications^16^. Indeed, immunocytochemistry confirmed the preference of kinesin-1 for acetylated microtubules in COS-7 cells (Fig. 3e). Consistent with our trajectory analysis, these acetylated microtubules are highly curved (Fig. 3e and Supplementary Fig. 3b). To test whether these microtubules were also fluctuating more, we imaged fluorescent microtubules together with kinesin-1, which revealed that microtubules decorated by kinesin-1 changed their shape over time by undulation-like motion (Fig. 3f-g and Supplementary Movie 4). This type of motion has been observed previously in *in vitro* microtubule gliding assays where immobilized microtubule motors propel partially immobilized microtubules^21^ and *in vivo* in *Xenopus* melanophores^22^ and epithelial cells^23,24^. These results suggest that the subset of microtubules labeled by kinesin-1 undergoes constant undulations under the action of some localized force generators that, like kinesin-1, preferentially interact with this microtubule subset.

### Microtubule bending fluctuations are suppressed by actin

Previous work suggested that microtubule bending is caused by actomyosin contractility^11^. To quantify undulating microtubule movement and the contribution of the actin cytoskeleton to it, we imaged live COS-7 cells transfected with mCherry-tubulin using spinning disk microscopy (Fig.4a and Supplementary Movie 5). The dynamics of lateral displacements of microtubules can be easily visualized using kymographs built along lines parallel to the periphery of cells (Fig.4b). To explore the influence of the embedding actin meshwork on the generation of these fluctuations, we treated cells with drugs having different effects: 10 μM of latrunculin A to depolymerize actin meshwork, 10 μM jasplakinolide to promote F-actin stabilization and polymerization and 50 μM blebbistatin to inhibit the contractility of myosin II motors. We quantified their effect on the lateral microtubule fluctuations using spatiotemporal image correlation spectroscopy ^25^. iMSD analysis revealed (Fig.4c-e) that myosin inhibition did not abolish fluctuations, while the F-actin-disrupting drug latrunculin A resulted in a twofold increase of both the speed (diffusion coefficient, Fig.4d) and confinement size of fluctuations (Fig.4e). Thus, microtubule fluctuations are not caused by the contracting actin network, but are instead dampened by it.

**Figure 4:**
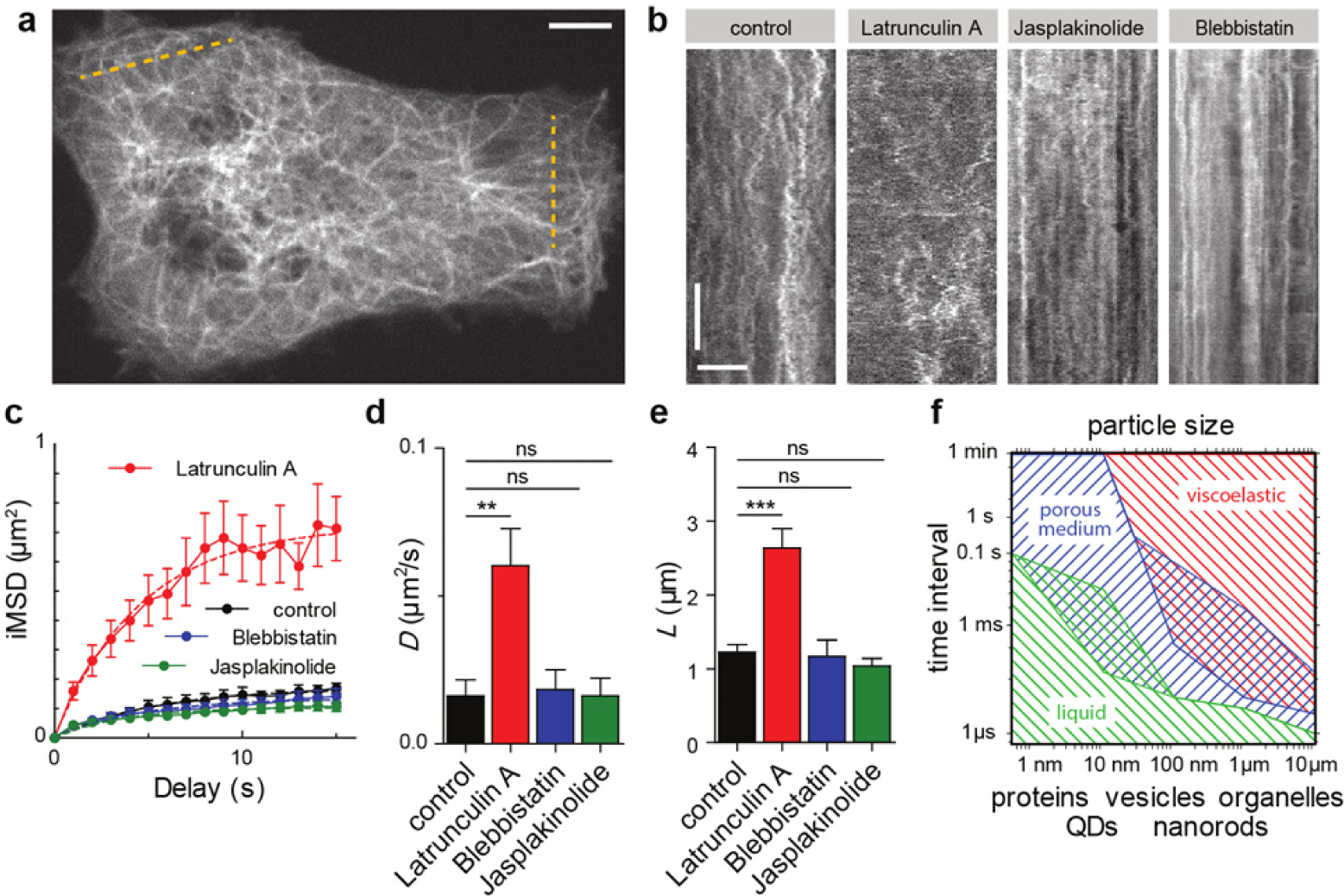
Analysis of microtubules bending deformations under the treatment of F-actin modifying drugs. (a) Still from a spinning disk movie of COS-7 cells expressing mCherry-tubulin. Dashed yellow lines mark areas used to build kymographs. Scale bar is 5 μm. (b) Representative kymographs built from spinning disk movies of COS-7 cells transfected with mCherry-tubulin under the treatment of indicated drugs. Scale bars are 60 s and 5 μm. (c) Plot of average iMSD versus time derived from kymograph analysis (N=11, 10, 14 and 13 for control, 10 μM latrunculin A, 10 μM jasplakinolide and 50 μM blebbistatin treatment). Error bars represent SEM. (d) Average diffusion coefficient derived from individual fitting of iMSD curves for indicated conditions. Error bars represent SEM. (e) Average confinement size derived from individual fitting of iMSD curves for indicated conditions. Error bars represent SEM. (f) Hypothetical phase diagram reflecting the behavior of intracellular particles and cellular components of different sizes on different time scales.

## Discussion

Based on the very slow diffusion of beads and SWNTs observed in earlier works^10,11^, the cytoplasm has recently been proposed to be a dense elastic network in which most particles are trapped in the actin meshwork. In this situation most diffusion-like behavior would be established by active contractions and remodeling of the actin network and therefore can emerge only at longer times scales^10,11^. By fast tracking of non-functionalized QDs, we instead revealed two populations of diffusive QDs that differed in diffusion constant by almost two orders of magnitude and the faster fraction could be increased by F-actin depolymerization. The fast population has a diffusion constant that is surprisingly close to the diffusion constant expected for these QDs in water (*D_fast_*~10 versus *D_water_*~16 μm^2^ s^−1^, Supplementary Fig.1a). It is likely that this fraction has been overlooked in many earlier experiments, because slower acquisitions will not detect this population and be biased towards the slower diffusing QDs embedded in actin meshwork. Detecting this subpopulation is also challenging using bulk average methods such as FRAP and FCS. The measured diffusion coefficient values allow us to estimate the viscosity of the aqueous phase of cytoplasm being 1.6 times higher than that of water, which is in agreement with previous estimates using bulk average techniques with small fluorescent molecules^26^.

Our results support a unifying model in which the slow diffusion observed here and in earlier work reflects the dynamics in specific actin-rich cellular subdomains, such as the actin cortex. The simultaneous co-presence of fast diffusion is attributed to a different, central sub-cellular compartment containing a less dense filament network. In addition, our results highlight the influence of probe size and geometry in probing cell environment. From super-resolution studies it is known that actin in the lamella of COS-7 cells forms two dense horizontal layers of 30-50 nm mesh size separated vertically by approximately 100 nm^15^. Particles of 100-300 nm in size will be trapped between these layers and their dynamics will reflect the fluctuations or remodeling of the network itself. For probe dimensions smaller than the mesh size, the assumption of a continuous viscoelastic cytoplasm is no longer valid and actin filaments will appear as discrete mechanical obstacles forming void compartments (pores) accessible for diffusion^7,8^ (Fig.4e). The ratio of *D_slow_*/*D_fast_* ~ 0.006 allows us to estimate the average pore size to be 36 nm (^27^ and Supplementary material), i.e. compatible with the size of the particle itself (~28 nm). Such probes will diffuse freely and rapidly inside the actin pores on time scales below (*d*_mesh_-*d*_probe_)^2^/6*D* ≈ (0.036-0.028)^2^/10 ≈ 5 μs, but will jump from one cell of the mesh to another at much longer timescales, resulting in a much slower effective diffusion constant at higher timescales (Fig. 1f).

What implications does this finding have for the intracellular transport of cargoes driven by different kinesin motors? To study this, we probed the active linear motion of QDs labelled with different types of kinesin motors. We developed a robust method to separate transient periods of directional movement from the random movement that interspersed these episodes and also carefully examined how longitudinal motility and transverse fluctuations should be extracted from the *xy*-trajectories. We found that large apparent transverse fluctuations emerge if the microtubule curvature is underestimated, but are absent if the microtubule curvature is closely followed by smooth fitted B-spline. This latter approach is preferable, because motors typically travel along a curve within 1-3 seconds, which is faster than the typical curvature remodeling time that we observed. In our study we therefore did not observe the large transverse displacements (~0.5-1μm) reported earlier for kinesin-1 fused to nanorods of 300-1000 nm long^11^. However, we did detect that the trajectories of this motor were both less smooth and more curved compared to other kinesins, which originated from its preferential binding to a subset of microtubules that underwent continuous bending and shaking. Preferential binding of kinesin-1 to modified microtubules has been previously observed, but the precise molecular origin of this selectivity has remained elusive^16^. Interestingly, we found that these preferred microtubules also undergo more active shape remodeling, again reflecting the ability of these microtubules to attract specific force generators.

Our experiments with actin-altering drugs provide evidence that the undulations of microtubules is not caused by contraction of the actomyosin network as suggested earlier^11^, but instead is strongly suppressed by it. These observations are in line with previous studies, showing that bending of microtubules is caused by microtubule specific motor generators^22-24^. To illustrate our findings and aid future studies of intracellular transport and cytosol compartmentalization, we built a hypothetical “phase diagram” of the heterogeneous cell environment (Fig.4f). The diagram maps the boundaries of applicable physical models of the cytoplasm as a function of probe sizes and time scales. For example, for proteins of 2-6 nm in size the cytoplasm can be considered as a liquid on almost any time scale. The behavior of bigger objects (20-100 nm) would depend on the time scale, but also on the position within the cell. Areas of overlapping hatching highlight “metastable” conditions where two compartments with different properties coexist simultaneously, as shown here for our QDs probes (time scale: 1 ms, size: 30 nm). Objects of 300-500 nm is size would be mostly stuck inside the filament network and move together with it. Further development of scalable tracers should allow precise mapping of those boundaries and a thorough description of the non-equilibrium mechanical environment of the cell^28^.

In summary, using novel functionalization, delivery and analysis tools, we found that the heterogeneity of the mammalian cytoskeleton imposes a large spatial variation in non-equilibrium cellular dynamics, which precludes straightforward application of physical approaches that model the cytoplasm as a viscoelastic homogeneous and isotropic medium. These results increase our understanding of the material properties of the cytoplasm, can guide future modeling approaches and could aid studies of passive delivery of nanoparticles or therapeutic agents.

## Methods

Live and fixed cells imaging, immunocytochemistry, GFP and nanobody functionalized QD stoichiometry analysis are described in Supplementary Methods.

### Cell culture and transfections

COS-7 were cultured at 37°C in DMEM/Ham’s F10 (50/50%) medium supplemented with 10% FCS and 1% penicillin/streptomycin. One to three days before transfection, cells were plated on 19 or 24 mm diameter glass. Cells were transfected with Fugene6 transfection reagent (Promega) according to the manufacturers protocol and grown for 16-24 hours. Human HA-KIF5B(1-807)-GFP-FRB, human KIF17(1-547)-GFP-FRB, rat KIF1A(1-383) and rat KIF21B(1-415) plasmid constructs were used for transfections, see Supplementary Methods for these and additional cDNA constructs information.

### Purification of recombinant nanobody

Bio-VHH_GFP_ was cloned in pMXB10 vector using vhhGFP4 sequence^29^. Recombinant bacterially expressed bio-VHH_GFP_ or bio-FKBP(2x) were obtained by using IMPACT Intein purification system. Induction, expression and purification of fusion proteins were performed according to the manufacturer’s instructions (New England Biolabs), see Supplementary Methods for details.

### Electroporation of COS-7 cells and functionalization of QDs

For electroporation of adherent COS-7 cells (all values are given for one 24 mm coverslip), 2 μl of Qdot 625 streptavidin conjugate (1 μM; A10196, Molecular Probes, Life sciences) and 20-25 μl of purified bio-VHH_GFP_ or bio-FKBP(2x) (0.7-0.8 μg/μl) were diluted in PBS to a final volume of 200 μl. Reaction was incubated for 1 hour at room temperature and then at 4°C overnight. Cells were electroporated with the Nepa21 Electroporation system (Nepagene). Electroporation was performed in 6-well plate containing 1.8 ml of warm Ringer’s solution (10 mM Hepes, 155 mM NaCl, 1 mM CaCl_2_, 1 mM MgCl_2_, 2 mM NaH_2_PO_4_, 10 mM glucose, pH 7.2) and 200 μl of electroporation mix. Parameters for electroporation (Voltage, Interval, Decay, Number and Pulse Length) were optimized from standard settings to achieve optimal efficiency (Supplementary Table 1). Each coverslip was electroporated with fresh solution of QDs. Cells were then washed with Ringer’s solution and either mounted in imaging ring for live imaging experiments or returned back to the growth medium and fixed at different time points.

### Particle detection, tracking and trajectories analysis

Image/movies processing routines were automated using ImageJ/FIJI macros or custom build plugins. MSD calculation, curve fitting and all other statistical and numerical data analysis were performed in Matlab (MATLAB R2011b; MathWorks) and GraphPad Prism (ver.5.02, GraphPad Software). Briefly, positions of individual QDs were determined by fitting elliptical Gaussian, linked to trajectories using nearest neighbor algorithm, manually inspected and corrected. Only tracks longer than 10 frames were used for analysis. MSD and velocity autocorrelation curves together with diffusion coefficient calculations were performed using “msdanalyzer” Matlab class^30^. Ensemble diffusion coefficients were measured as a slope of the affine regression line fitted to the first 25% of weighted average MSD curves and divided by four (assuming two dimensional motion). Detection of motor runs and spline fitting of kinesins’ trajectories are described in Supplementary Methods.

## Acknowledgments

This research was supported by the Dutch Technology Foundation STW and the Foundation for Fundamental Research on Matter (FOM), which are part of the Netherlands Organisation for Scientific Research (NWO). Additional support came from NWO (NWO-ALW-VICI to C.C.H and NWO-ALW-VIDI to L.C.K.) and the European Research Council (ERC Starting Grant to L.C.K.). M.M. is recipient of a European Molecular Biology Organization (EMBO) Long-Term Fellowship (EMBO ALTF 884-2011), Marie Curie IEF (FP7-PEOPLE-2011-IEF) and Deutsche Forschungsgemeinschaft Emmy-Noether Programm (Ml 1923/1-1). We thank Dominique Thies-Weesie for help with viscosity measurements and Boris Slepchenko for comments on pore size estimation.

## Author Contributions

E.A.K., M.M., A.A., C.C.H. and L.C.K. designed the experiments. M.M., E.A.K. and H.X.B. performed experiments. E.A.K. and M.M. analysed the data. P.B.H. contributed reagents and experimental suggestions. All authors discussed the data and commented on the manuscript. E.A.K., M.M. and L.C.K wrote the paper. L.C.K supervised the project.

